# MCA: A Multicellular analysis Calcium Imaging toolbox for ImageJ

**DOI:** 10.1101/2025.08.19.671108

**Authors:** John Hageter, Audrey DelGaudio, Maegan Leathery, Braxton Johnson, Tegan Raupp, James Holcomb, Axel Faz Treviño, Julius Jonaitis, Morgan S. Bridi, Andrew Dacks, Eric J. Horstick

## Abstract

Functional imaging using genetically encoded indicators, such as GCaMP, has become a foundational tool for in vivo experiments and allows for the analysis of cellular dynamics, sensory processing, and cellular communication. However, large scale or complex functional imaging experiments pose analytical challenges. Many programs have worked to create pipelines to address these challenges, however, most platforms require proprietary software, impose operational restrictions, offer limited outputs, or require significant knowledge of various programming languages, which collectively can limit utility. To address this, we designed MCA (a Multicellular Analysis toolkit) to work with ImageJ, a widely used open-source software which has been the standard image analysis platform for the last 30 years. We developed MCA to be visually intuitive, utilizing ImageJ’s platform to generate new images based on completed tasks so users can visually see each step in the analysis pipeline. In addition, MCA implements a user-friendly GUI providing a simple interface which resembles other native ImageJ plugins. We incorporated functionality for rigid registration to correct motion artifacts, algorithms for cell body prediction, and methods for annotating cells and exporting data. For cell prediction, we trained a custom model in Cellpose 2.0 for segmentation of nuclei expressing pan-neuronal nuclear localized GCaMP in zebrafish. We validated the accuracy of MCA output to previously published zebrafish calcium imaging data which elicited visually evoked neuronal responses. To show the versatility of MCA, we also show that our software can be utilized for multiple sensory modalities, brain regions, and multiple model organisms including *Drosophila* and mouse. Together these data show that MCA is viable for extracting calcium dynamics in a user-friendly environment for multiple forms of functional imaging.

**Motivation:** Calcium imaging has become one of the most common methods for investigating neural activity, however analytical methods are limited to a few software platforms or are custom made. This limits replicability and imposes restrictions on incorporating additional tools to support analysis. To address these challenges, we developed a modular, graphical based, open-source toolbox, based in the ImageJ application, for performing functional imaging analysis in diverse models and datasets.

**Highlights:**

1. **Developed MCA, an ImageJ based plugin for analyzing functional imaging datasets.**
2. **Validated accuracy of MCA functions**
3. **Utilized MCA across multiple sensory modalities and model organisms**

## Introduction

Functional imaging using genetically encoded indicators (GEIs) has become an invaluable and widely used strategy to assess cellular function. Arguably, the most prominent and widely used GEIs are genetically encoded calcium indicators, such as GCaMP, a fluorescent reporter of calcium concentrations and proxy for cellular activity.^1–3^ However, calcium is not the only relevant GEI. GEI tools are available for glutamate, GABA, voltage, glucose, dopamine, and many other key biological factors.^2,4–6^ Availability of such tools has given researchers an unprecedented ability to assess the function of nearly every cell type and organ system in the body, often with cellular resolution and in vivo. The utility of GEIs is evident by the rapidly growing evolution of these tools to fluorescently observe and evaluate diverse neurotransmitters, cell types, or regions.^4,5,7^ The ability to leverage genetic control has also opened the ability to localize indicators with genetic tags. For example, nuclear localization, synaptic localization, or other tagged localization of GCaMP have allowed for improved single cell segmentation or synapse characterization in neurons, respectively.^4,8^ Moreover, the advancement of GEIs in combination with ever improving imaging technologies, like multiphoton and light sheet imaging strategies, has allowed functional characterization to expand to whole tissue, multi-regional brain planes, and even whole brains. Therefore, GEIs are capable of cell to system scale characterization, which is viable across numerous species.^6,9–11^ However, despite the widespread usage and application of functional imaging with GEIs, a major challenge is to perform analysis that extracts meaningful, quantifiable data from datasets containing hundreds if not thousands of cells. For instance, extracting cell or region-specific signals from large-scale imaging sets, sample-to-sample registration for rigorous spatial comparison, and predicting neuronal locations is non-trivial and can readily become computationally complex or excessively time-consuming if performed manually. Therefore, many functional imaging analyses require software specifically developed for the task. Indeed, several programs have been developed, yet in most cases require optimization through command-line inputs or access to proprietary software.^12–15^ An alternative is that many labs develop custom code^7,14,16^, yet these may not be readily adopted by other labs depending on programming skill or available hardware or software. Therefore, a gap exists between the accessibility of GEI tools for functional imaging acquisition and ability to readily and rigorously extract this data.

To address this gap we developed MCA, a <u>M</u>ulti-<u>C</u>ellular <u>A</u>nalysis toolkit developed in the Java programing language for use with ImageJ, a completely open-source and widely used scientific image analysis program^17^. MCA’s primary software tool, the Cell Manager plugin, mimics the functionality of ImageJ’s built-in ROI Manager providing a simple user interface with methods designed specifically for functional imaging analysis by extending the functionality of plugins already available in ImageJ as well as providing new functionality not currently available. MCA provides a simple graphical user interface (GUI), executed within ImageJ, and will perform image registration for motion correction, single cell segmentation, custom ROI based annotation, data standardization based on user selected parameters, averaging across replicates, background subtraction, and writing of functional imaging data from every form of image type available for use in ImageJ. Moreover, MCA includes spreadsheet format output for simplified analysis in programs such as Microsoft Office or Libre Office. We validated MCA using GCaMP imaging in larval zebrafish in response to visual and auditory stimuli spanning several brain regions. In addition, we leverage GCaMP imaging datasets from *Drosophila melanogaster* and *Mus musculus* to show that MCA is tractable for multiple species analyses and detection of cellular or regional activity. Collectively, we demonstrate that MCA provides a straight-forward analysis application for complex functional imaging data that is tractable and integrated into a widely used and open-source image analysis software.

## Results

### <u>M</u>ulti-<u>C</u>ellular <u>A</u>nalysis toolkit is a GUI plugin for ImageJ to analyze functional imaging datasets

We designed a new functional imaging software for ImageJ for two main reasons. 1) ImageJ is one of the most widely used tools for scientific imaging analysis while remaining open source and freely available to the public^17,18^, and 2) many of the current functional imaging packages available pose specific challenges such as steep learning curves or relying on proprietary software impeding accessibility. We propose that integration with ImageJ would provide access to diverse and powerful resources for image analysis, especially if coupled with functional imaging capabilities. Thousands of plugins have been developed for ImageJ that have provided expansive resources for image analysis.^19–21^ However, what remains lacking is a centralized, flexible, and user-friendly functional imaging analysis plugin for ImageJ that could leverage the currently available applications associated with this widely used resource.

The main interface of the MCA plugin is the cell manager GUI. The GUI consists of two primary sections: 1) a list of regions of interest (ROIs) that contain the data for each ROI and 2) a list of buttons that facilitate key processes in functional imaging analysis (Figure 1A). In addition, we added two tabs for organizing ROIs where the user can look at individual ROIs in the “Cells” tab or user-defined groups of cells in the “Groups” tab. GUI functional buttons assist in key processes of calcium imaging analysis. These functions are designed to be modular, meaning they can be used in any order or with any other available ImageJ function (Figure 1B). Each one of the functions listed are designed to provide visual user feedback with output that shows how the function processed the image (Supplemental Figure 1A). The goal of MCA’s architecture is to allow researchers to visually understand how their data is being processed and restrictions or processes happening “behind the scenes” throughout analytical steps. In addition, this is software is supported by the ImageJ Bio-Formats plugin that allows for the conversion of most proprietary imaging formats to ImageJ’s ImagePlus format used by MCA (Figure 1C).^22^

**Figure 1:**
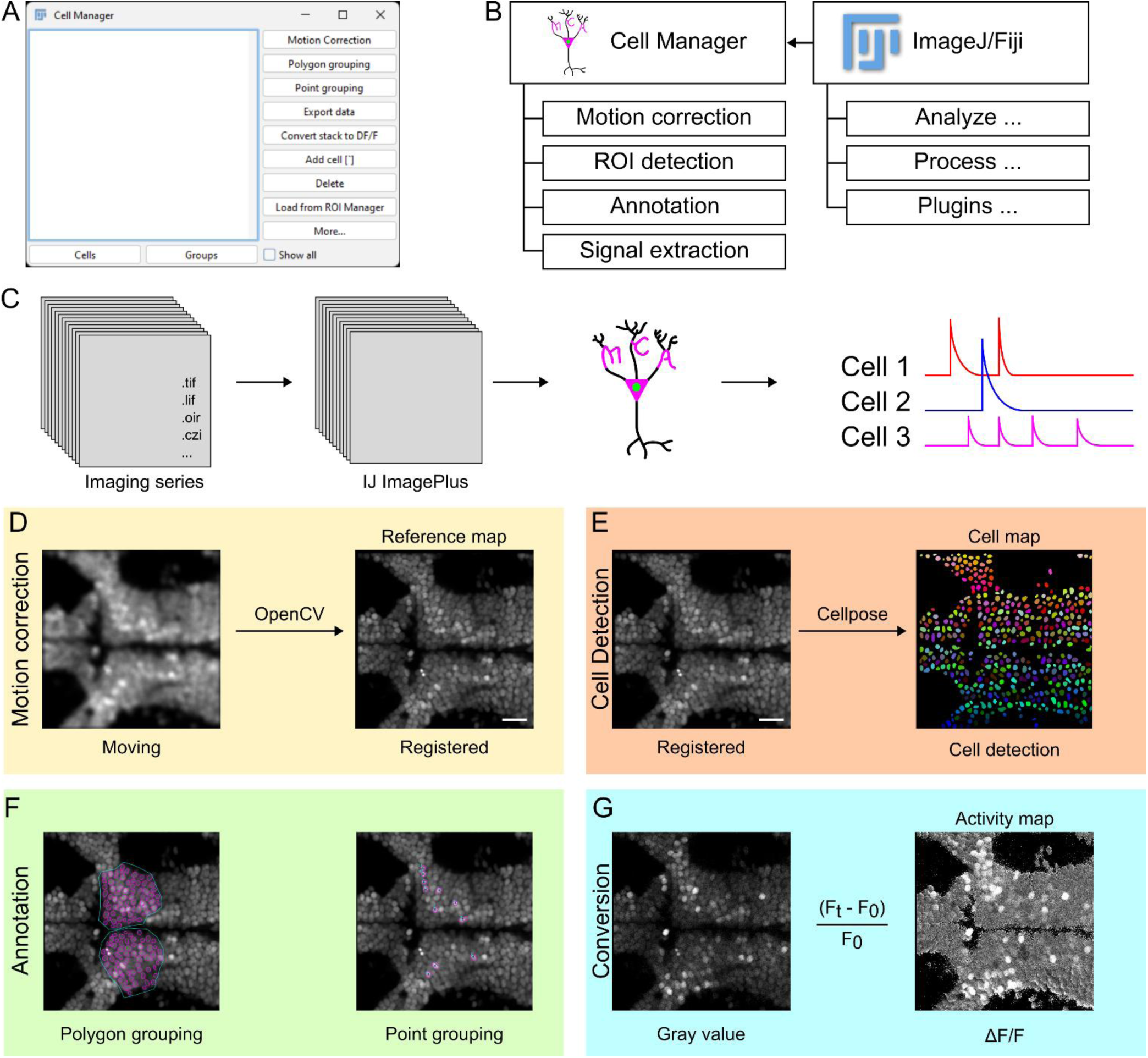
MCA is a calcium imaging toolbox integrated with ImageJ. **A.** Screenshot of the Cell Manager GUI and functional layout. **B.** Functions available in the Cell Manager and representative functions that can integrate with the Cell Manager from Fiji/ImageJ. **C.** Representative import of most imaging formats to the ImageJ ImagePlus format which is used for all analysis in MCA. **D.** Representative of motion correction used by MCA integrating the OpenCV template matching algorithm. **E.** Representative cell segmentation of a sum projection of an imaging series using a trained H2B-GCaMP model for cell detection in Cellpose. **F.** Representatives for polygonal grouping of cells or point grouping. **G.** Illustrative example of imaging series converted from an acquired 8-bit gray value image into a 32-bit image series that represents the ΔF/F values at each pixel.

In the Cell Manager, we integrated processing functions for motion correction, ROI detection, cellular annotation, and signal extraction for analyzing data, and lastly a function which exports data from ImageJ to spreadsheet-based format. Typically, functional imaging videos are subject to motion artifacts from animals moving, minor vibrations from imaging equipment, or external factors such as human disruption. To correct this, we implemented OpenCV’s template matching algorithm, which is the main method used in the “Template Matching” plugin developed for ImageJ (Figure 1D)^23,24^. This is an intensity based method for correcting motion artifacts, which has been widely used for simple rigid motion correction in calcium imaging and other time series imaging.^24–26^ A main function of the Cell Manager is the ROI detection. The goal for this method was to integrate an approach which automates cell detection and segmentation from an image. To achieve this, we incorporated two machine learning model-based approaches for detecting cells from an image; Cellpose and StarDist.^27–30^ These tools are machine learning models that are regularly used to detect cell boundaries and designed to aid in segmentation of cell nuclei (StarDist) or detect different types of cells (Cellpose). Both of these model based approaches have been utilized for cell segmentation of biologic data^31–34^ and advanced biologic image processing applications.^35–37^ When used in MCA, Cellpose will return an image of predicted cell locations from an input image while StarDist imports cell ROIs directly into ImageJ’s ROI manager (Figure 1E). From either of these outputs, cell ROIs can be directly imported from the ROI manager to the Cell Manager with the “Load from ROI Manager” which syncs ROI data between each manager. To maintain MCA as a user-friendly plugin, we integrated the capacity for annotation of data prior to exporting data. These methods facilitate quick renaming of cellular ROIs and the ability to group ROIs together. The main annotation methods are grouping functions which allow users to group cell ROIs together using the reference map by either manually drawing a grouping ROI around multiple cells with the “Polygon grouping” function, or by selecting multiple cells individually with the “Point grouping" function (Figure 1F). This method of annotating groups of cells together prepares them for export by specific user defined groups providing easy organization of large imaging areas with cells that may belong in unique categories.

Next, we wanted to design a clear way to make raw data immediately accessible and understandable through visualization, reducing the need for users to explore data through typical file formats such as binary or CSVs. To achieve this, we sought to convert an imaging series from the original pixel values to values that would be directly used when exporting data. Standard image pixel values are not useful in measuring activity because they vary wildly among samples and are sensitive to GEI expression levels. Through the “Convert stack to DF/F” function, pixel values from the recording are converted to standardized change in fluorescence (ΔF/F) relative to a user defined baseline period, which is calculated for every pixel (Figure 1G). This transformation standardizes fluorescence intensity values, among cells and across individuals, in a visual manner rather than spreadsheet-based format.

Data also typically needs to be filtered to reduce noise, which supports identifying where biologically relevant activity is occurring. MCA accomplishes this through two distinct functions that can be applied when exporting data: 1) signal filtering, and 2) peak detection. Currently, MCA uses a gaussian filter, which smooths fluctuations across data points in the raw fluorescent signal. The algorithm for detecting peaking activity is a sliding window-based method, which has been widely used in a variety of applications requiring accurate estimation of changes during time-series data.^38–42^ This method looks for sudden changes or “peaks” in the calcium signal over time using a user-defined time window and comparing signal change across preceding windows. The threshold for significance is based on the number of standard deviation changes, which is also user-defined. This peak detection algorithm indicates within the recording where there was no change (value=0), significant increase (value=1), or significant decrease (value=-1) in signal activity. We reasoned that this peak detection output would provide a simple interface for further analysis and data exploration. Data is then exported from MCA through the “Export Data” function. This function compiles data from all the ROIs in the Cell Manager, group annotations, and image stacks from multiple recordings and exports to two csv files, which represent signal data (ΔF/F format) and predicted significant peaking activity (0, 1, -1 format). Exported data will contain columns for every slice of the image stack and the corresponding signal value. Additional columns denote the name of the file recorded from (“Name” column), the name of the ROI (“ROI” column), X and Y coordinates for the center point of the ROI (“X” and “Y” columns), as well as the method used for filtering (“Filter” column) and detecting peaking activity (“Detection.Method” column) from the signal data.

MCA can be leveraged in many ways to process functional imaging data, which we outline a generic workflow for using MCA (Figure 2A). Broadly, there are two paths which can be flexibly used and adjusted based on experimental goals that we refer to as the ‘analysis’ and ‘annotation’ paths. The analysis path encompasses the steps required for exporting data and includes motion correction, cell labeling (either manual or using a model), converting the stack to ΔF/F, and exporting data to csv format. This path can, at any step, be accompanied by the annotation path which includes functions for organizing and labeling ROIs which eases post-data export analyses. Generating a “map” is not a function contained within MCA but can be accomplished with basic ImageJ functions. For example, using the reference map, ROIs can be easily labeled or grouped using the available methods. After grouping cells together, these groups and the ROIs contained within will be available to visualize from the “Groups” tab in the GUI or in a dedicated “Groups” column within the exported csv. Together, these features and architectural design make MCA a comprehensive and user-friendly plugin for working with functional imaging data in ImageJ.

**Figure 2:**
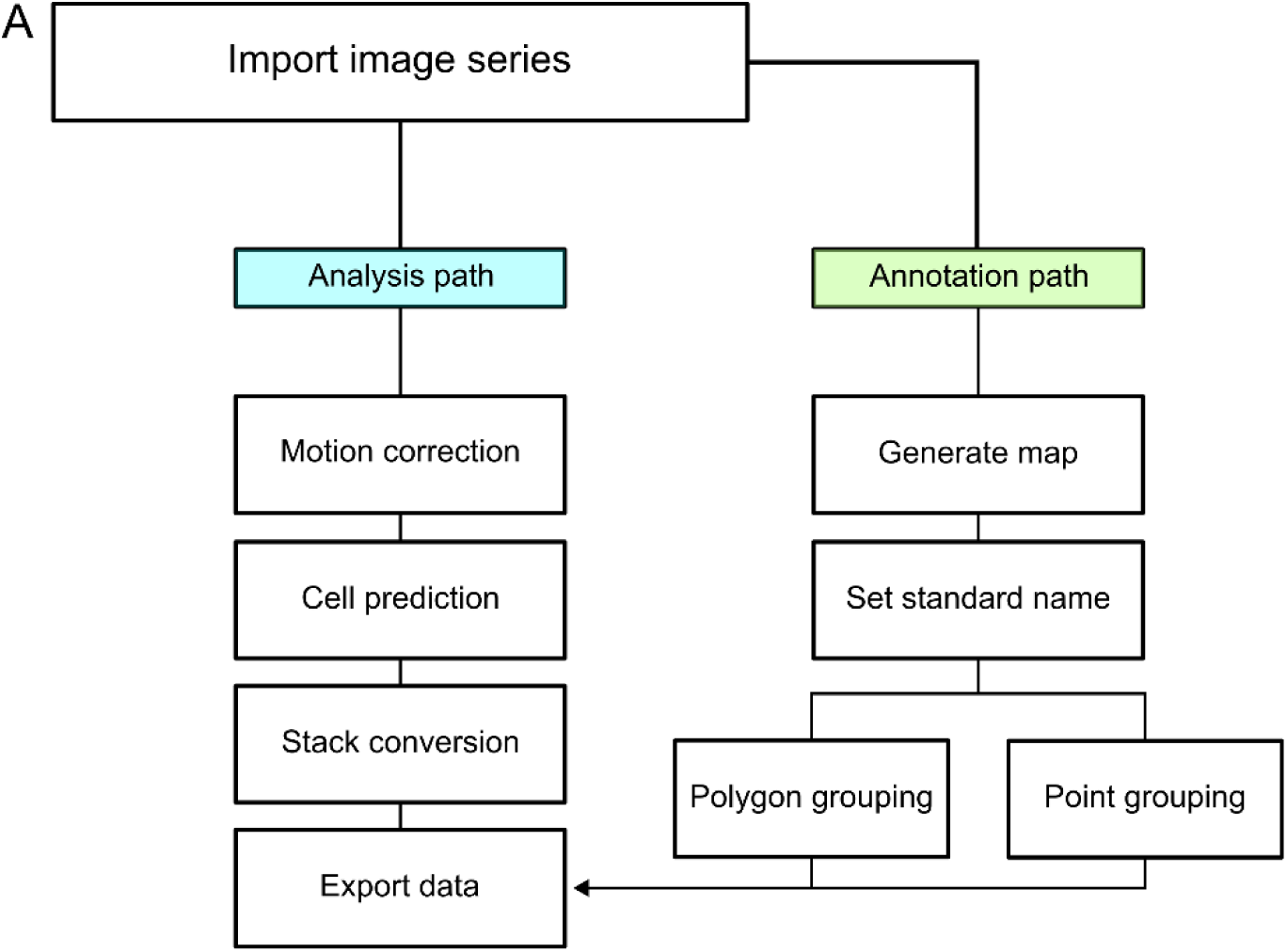
Workflow for using MCA. **A.** Representative workflow following the import of an imaging series into ImageJ. There are two main paths for working in the Cell Manager. An analytical path and an annotation path. The annotation path consists of using external functions, like creating a composite image or max projections to create a map which can be annotated. From here the user can set standard naming to ROIs and then group ROIs with two methods, polygonal grouping or point based grouping. The analysis path is required to generate a dataset from the imaging series. Initially begin with the motion correction step, continue to the cell detection step by using either Cellpose or Stardist2d to predict cellular ROIs, or manually label them. Next, the imaging series is converted to raw data and finally that data is exported into spreadsheets.

### MCA is validated against manually analyzed data

To validate the accuracy of MCA, we repeated an analysis of a previously published calcium imaging dataset. In the previous analysis visual stimulation evoked distinct patterns of neuronal activity across thalamic regions and across brain hemispheres.^31,44^ We reasoned these differential responses would provide an ideal dataset to test the ability of MCA to extract biologically relevant significance. This experiment used a pan-neuronal nuclear localized GCaMP line in-crossed to reporter lines, *Tg(elavl3:H2B-GCaMP6f; y279:Gal4; UAS:epNTR-tagRFPt)*, to visualize thalamic neurons of interest and surrounding unlabeled neuronal populations. Therefore, this analysis provided regionally distinct functional differences to detect, and neuron and brain region level segmentation for which to test MCA. Initially we wanted to determine how accurately our Cellpose model detected cellular ROIs. To do this, we applied sum projections to all 19 larvae in our dataset and applied the H2BGCaMP model we trained in Cellpose (see methods). For comparison, we manually counted all cells across these 19 larvae. We found that the number of cells identified by our trained Cellpose H2BGCaMP model and manual counts were highly similar, which supports the efficacy of the cell segmentation used in MCA (Supplemental Figure 1A-B). For this experiment, larvae are exposed to a light off stimulus and we filtered neurons which responded to light extinction (Figure 3A). From our new analysis using MCA, we quantified neurons as being light responsive if a response was, 1) at least 3 standard deviations greater than the standard deviation of the 30 previous timepoints using our implemented peak detection algorithm and 2), if at least 3 consecutive timepoints had values above this threshold. Based on their responses to the light stimulus, we grouped neurons into OFF responsive neurons, ON responsive neurons, or OFF/ON responsive neurons according to their intensity change either following the loss of illumination, return of illumination, or both, respectively. Using MCA, we grouped neurons into the same regions as described in the previous publication: Greater Thalamus (Great Th.), Lateral Thalamus (Lat Th.), and Posterior Tuberculum (PT) along with grouping these regions by hemisphere and association with behavior (Figure 3B).^16,31,45^ The MCA analysis and Cellpose segmentation produced similar quantification and hemispheric asymmetric patterns as previously reported (Figure 3C-D, Supplemental Figure 3A-D). Using MCA, we were able to extract a total of 958 responsive neurons across 19 fish (OFF-responsive=644; ON-responsive=163; OFF/ON-responsive=151). This compares to our published analysis where we were able to extract 976 responsive neurons across the same 19 fish (OFF-responsive=718; ON-responsive=224; OFF/ON-responsive=34) (Figure 3E, Supplemental Figure 3E). In our previous analysis, we were able to visualize asymmetric strength in response to the loss of illumination across different regions of the Thalamus, yet not the PT. To compare our previous findings with the MCA analysis we averaged OFF-responsive neurons response for the 3 timepoints following the loss of illumination. Across all three major regions of interest (Great Th., Lat. Th., PT) we noted an increased response strength. For the Great Th., automated ROI detection using MCA replicated the prior observation (which was obtained by manual ROI placement) that responses in the hemisphere matched to the behavior have a larger response than those located in the opposed hemisphere (Figure 3F green box MCA: t(162)=4.94, p<0.0001; Prior data: t(141)=4.71, p<0.0001). We also recapitulate significance in lateral thalamic AMNs where responses are stronger in the hemisphere opposed to the motor behavior (Figure 3F orange box; MCA: t(78)=-3.86, p=0.0002; Prior data: t(183)=-2.22, p=0.028). Lastly, we show that no asymmetry was present in the PT, consistent with previous findings (Figure 3F purple box MCA: t(248)=0., 913, p=0.362; Prior data: t(316)=-0.885, p=0.377). This analysis shows that MCA was able to successfully delineate key ROIs, perform automated segmentation, and from this recapitulate functional and hemispheric differences previously identified by laborious manual processing of functional imaging data. One difference we observed was that the MCA responses showed higher ΔF/F, which was likely due to the background subtraction included with the MCA toolkit. This validation work shows that automated ROI detection using MCA reliably and accurately generates output consistent with prior observations using manual methods of functional imaging analysis.

**Figure 3:**
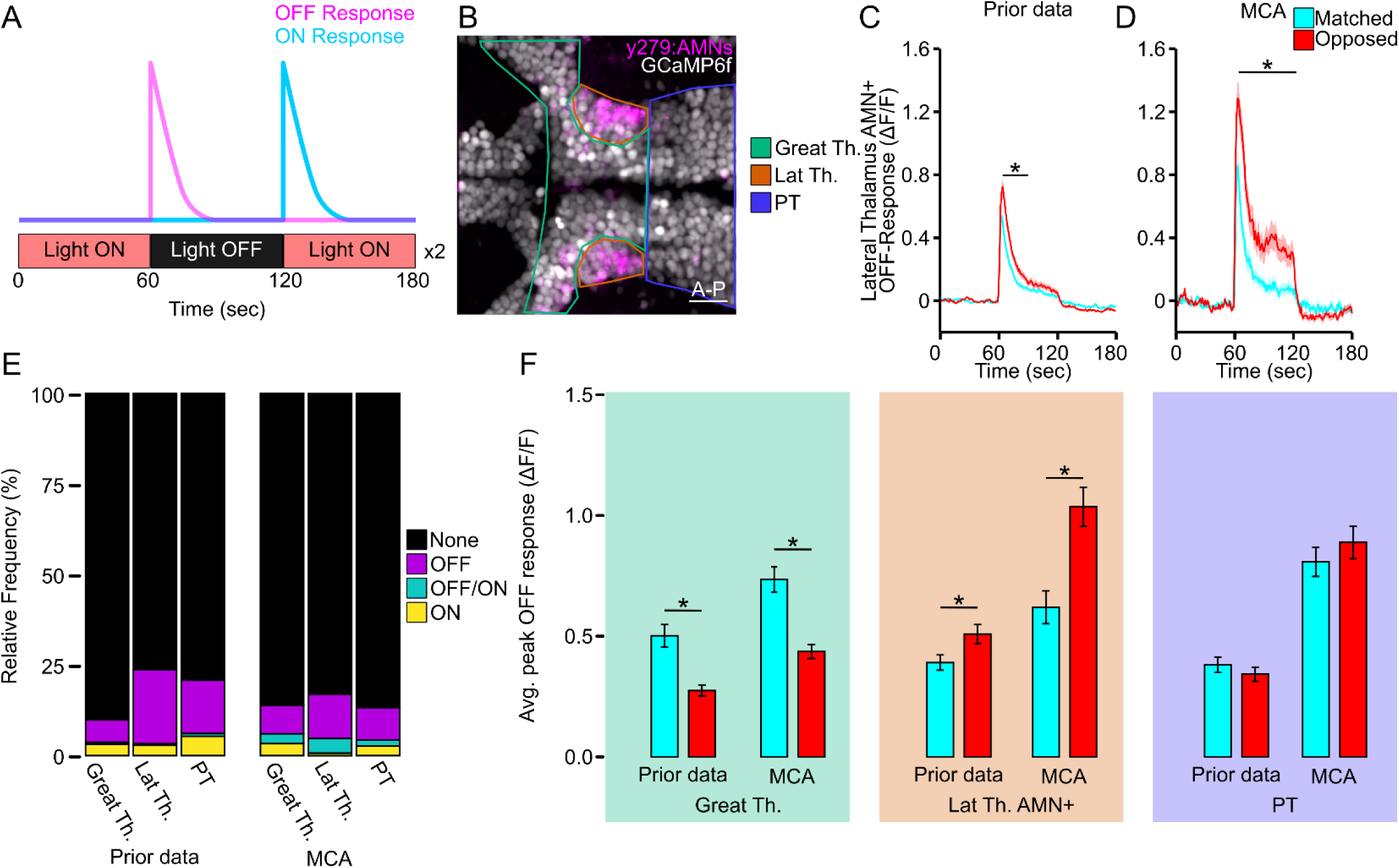
MCA maintains previous trends in calcium imaging data. **A.** Representative calcium imaging assay and representative OFF (magenta) or ON (cyan) response. Larvae are imaged for 3 minutes with the light on for 1 minute, light OFF for 1 minute, and light back on for 1 minute. This series is repeated twice, and data are averaged together. **B.** Representative sectioning of the Thalamus into the Lateral Thalamus (Lat Th., orange) which contains the AMNs (magenta), the Greater Thalamus (Great Th., green) which is every cell medial and anterior to the Lateral Thalamus, and the Posterior Tuberculum (PT, purple) scale bar 20µm. **C** Averaged output for OFF-responsive neurons in the AMNs between matched (cyan) and opposed (red) hemispheres for published data (Prior data: matched: N=79; opposed, N=106). * Indicates p < 0.05 between matched and opposed for at least 10 consecutive timepoints. **D.** Same as **C** yet using MCA (matched, N=50, opposed: N=60). * Indicates p<0.05 two tailed t-test between matched and opposed where p<0.05 for at least 20 consecutive timepoints. **E.** Relative frequency of response types among regions between analysis methods (Prior data: **Great Th.:** None (Black), N=1799 (83.2%); OFF (purple), N=143 (6.1%); OFF/ON (teal), N=13 (0.5%); ON (yellow), N=72 (3.2%); **Lat Th.**: None, N=939 (79.9%); OFF N=252 (20.5%); OFF/ON, N=5 (0.4%); ON, N=35 (2.8%); **PT**: None, N=1706 (79.2%); OFF N=318 (14.7%); OFF/ON, N=16 (0.7%); ON, N=115 (5.3%); MCA: **Great Th.:** None, N=1799 (86.1%); OFF, N=164 (7.9%); OFF/ON, N=57 (2.7%); ON, N=69 (3.3%); **Lat Th.**: None, N=606 (79.9%); OFF N=89 (12.2%); OFF/ON, N=30 (4.1%); ON, N=5 (0.7%); **PT**: None, N=2421 (86.8%); OFF N=250 (9.0%); OFF/ON, N=45 (1.6%); ON, N=74 (2.7%)). **F.** Average peak off response between matched (cyan) and opposed (red) OFF responsive neurons in the Greater Thalamus (green; Prior data: matched N=63, opposed N=80; MCA: matched N=84, opposed N=80), Lateral Thalamus AMN+ (orange; Prior data: matched N=79, opposed N=106; MCA: matched N=36, opposed N=44), and Posterior Tuberculum (purple; Prior data: matched N=170, opposed N=148) * indicates p<0.05 two-tailed t-test between matched and opposed within methods.

### MCA is versatile for a variety of experimental formats

Next, we wanted to demonstrate the flexibility of the MCA platform for analyzing functional imaging of sensory driven activity in the brain of other model organisms and sensory modalities. We first expanded our analysis by applying MCA to auditory responses in the larval zebrafish stato-acoustic ganglia (SAG). First, we designed a method to test acoustic responses from the SAG in zebrafish in response to an auditory stimulus. The SAG connects hair cells within the inner ear to the brain and are necessary for hearing and maintaining balance.^46–48^ We recorded a series of progressively increasing amplitude auditory cues to larval zebrafish utilizing a ventrally mounted speaker to deliver an acoustic stimulus (Figure 4A). We used the *Tg(y256:Gal4; UAS:epNTR-mCherry; elavl3:H2B-GCaMP6f*), which allowed visualization of the SAG and pan neuronal expression of nuclear localized GCaMP6f (Figure 4B). From this stimulation series, using MCA, we were able to extract auditory responses across all stimulus amplitudes (Figure 4C-D). We categorized significant cellular responses if at least two of five timepoints were significant for one or more frames following stimulus onset. We found that MCA was able to extract responses from 26.7% of SAG cells following an auditory stimulus (Figure 4E-F). Typically, only ∼2% of neurons in the zebrafish SAG respond to vibrational stimuli whereas 40% respond to changes in balance orientation.^49–51^ This is a larger percentage of responses compared to previous evidence, however, we posit this is due to the aggressive amplitudes used for delivering stimuli which could disrupt zebrafish balance sensation.

**Figure 4:**
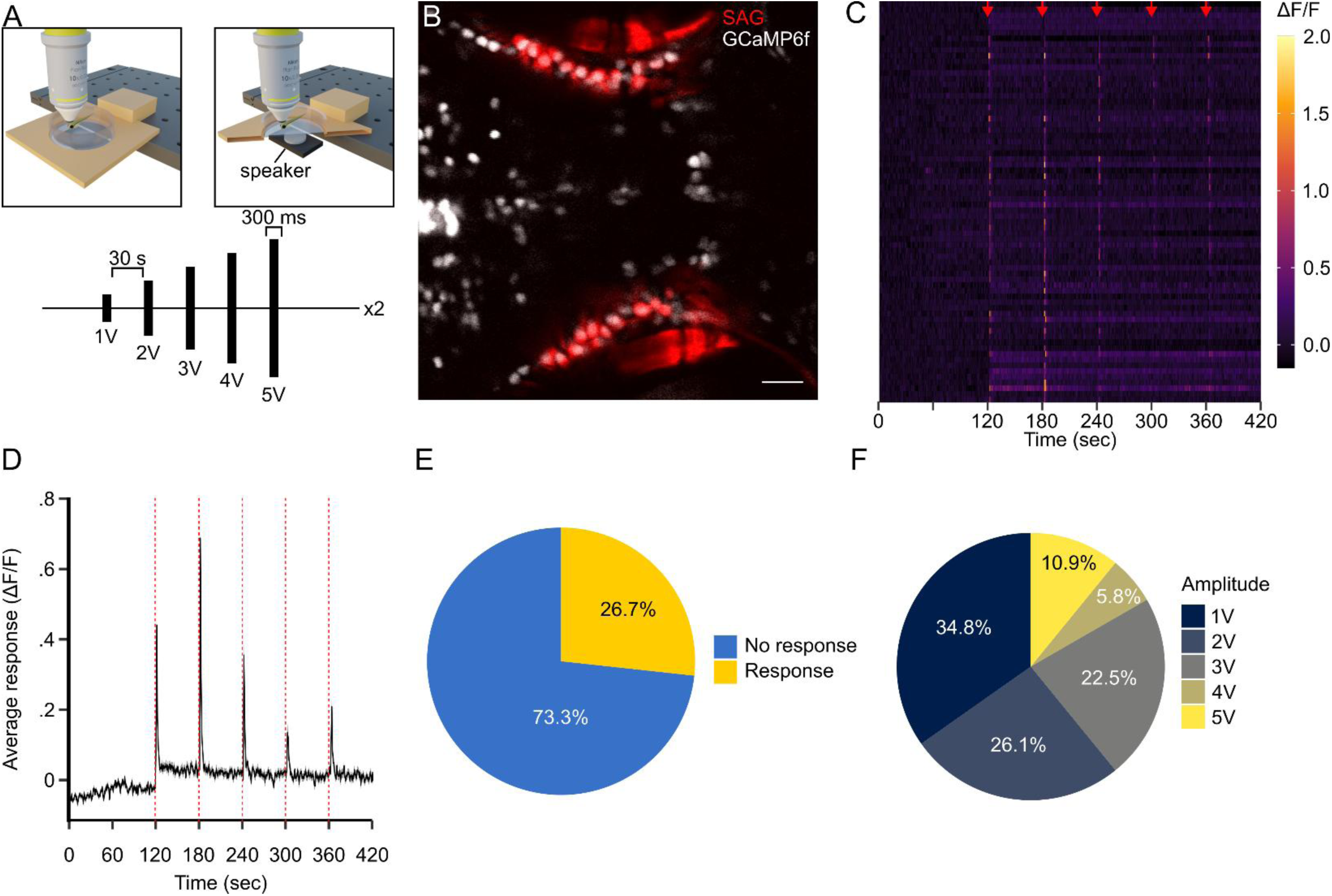
MCA is versatile for a variety of stimulus delivery techniques. **A.** Representative diagram of acoustic calcium imaging series where larvae are exposed to an acoustic stimulus of increasing amplitude every 30 seconds. **B.** Representative imaging plane indicating SAG cells (red) and pan-neuronal GCaMP6f (white). Scale bar (20 µm). **C.** Raster plot where every row is a different cell that responds to at least one of the acoustic stimuli (denoted by red arrows). Color indicates ΔF/F value. **D.** Average of all responsive cells through the imaging series. Line and ribbon indicate mean ± SEM. Dotted red lines indicate stimulus points. **E.** Pie chart indicating the percentage of SAG cells that responded to at least one of the acoustic stimuli (Response: gold, N=70; No Response: blue, N=192). **F.** Same as in **E** Indicating the percentage of responsive cells to each amplitude (1V, N=48; 2V, N=36; 3V, N=31; 4V, N=8; 5V, N=15)

To further demonstrate the utility of MCA, we wanted to show that this is a valuable toolkit for assessing functional imaging data from other model systems. Compared to our previous pipeline for extracting responses from nuclear localized GCaMP, our other model organism datasets used a cytosolic GCaMP. With ImageJ’s native ROI drawing tools, we were able to adjust our workflow to manually draw ROIs. We applied this adjusted MCA pipeline to a regional calcium imaging recording from the *Drosophila* antennal lobe where individual glomeruli were segmented to record responses from individual neuronal compartment to a timed exposure to apple cider vinegar (ACV), a common and potent olfactory stimulus (Figure 5A).^52,53^ The MCA pipeline was able to extract responses from local interneurons with electrotonically isolated compartments that innervate individual glomeruli (Figure 5B-C).^54^ We next applied MCA to a calcium imaging dataset from putative pyramidal cells in the mouse visual cortex. A visual stimulus of a grating pattern moving across a screen at randomized orientations was presented to test visual acuity.^55^ Using native ImageJ tools, we manually labeled cells within the imaging series and from MCA’s “Convert to DF/F” function was able to visualize neurons which had strong increases or decreases in activity during the recording (Figure 5D-E). MCA was then able to extract these cellular responses and when plotted in accordance with the stimulus presented, was able to easily visualize cells that increase in activity in response to the onset of the stimuli and those that decrease when the stimuli is removed (Figure 5F). Both analyses highlight the versatility of MCA to be applied to widely used experimental model systems and different sensory modalities.

**Figure 5:**
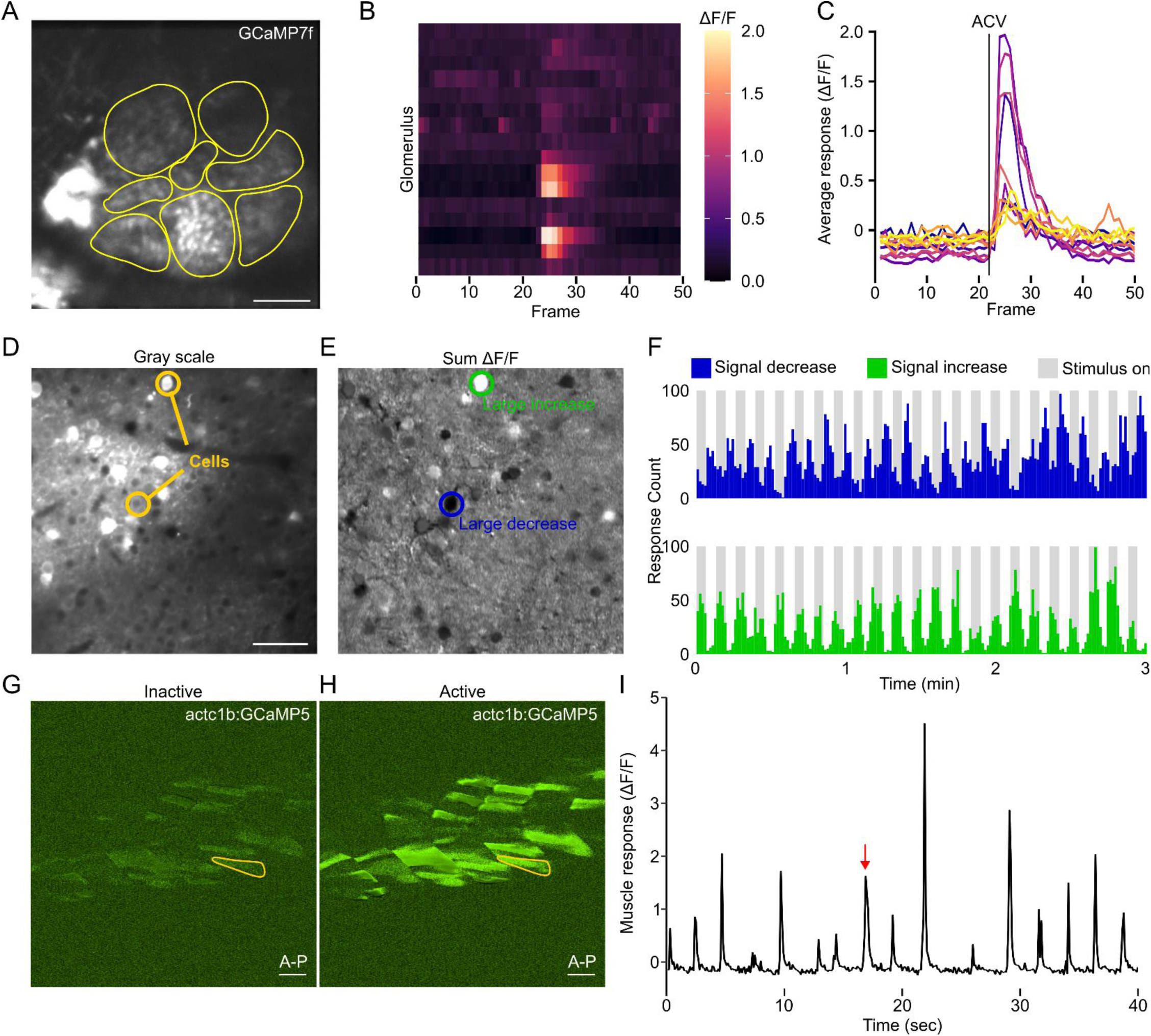
MCA is viable for multiple model organisms. **A.** Representative of imaging plane for recording odor evoked responses from local interneurons in the antennal lobe of *Drosophila* (scale bar=20µm). Yellow circles indicate locations of individual glomeruli. **B.** Raster plot where every row is a different glomerulus (N=16) and response to ACV stimulation at frame 23. Color indicates ΔF/F value. **C.** Average glomeruli response. **D.** Representative of mouse calcium imaging plane in V1 of the superficial visual cortex (scale bar=20µm). **E.** Activity map generated from recording use in **D.** Green outline highlights an example cell where cell signal was significantly increasing, while blue highlights significantly decreasing. **F.** Number of significant decreases (top, blue) or increases (bottom, green) in signal activity among all neurons measured (N=98). Grey bars indicate visual stimulus is being presented to the mouse. **G-H.** Representative image of a zebrafish tail **G.** prior to spontaneous movement or **H.** during spontaneous movement. Yellow outline indicates cell being plotted in **I.** **I.** Calcium signal for single muscle cell outline in yellow in **G-H.** Red arrow denotes timepoint shown in **H.**

Last, we wanted to demonstrate that MCA is also viable for extracting activity from non-neuronal tissue types. Calcium imaging in muscle cells is a common practice used to understand the cellular dynamics caused by genetic disorders, tissue damage, or following regeneration.^56–60^ To demonstrate this, we recorded spontaneous muscle contractions in the zebrafish tail during a 3 minute recording. Zebrafish engage in spontaneous bouts of movement even when embedded in agar.^6^ Using this approach, we were able to record from skeletal muscle cells along the tail using *Tg(actc1b:GCaMP5)*^61^ which is expressed skeletal muscle.^62,63^ We were able to visualize spontaneous calcium flux in skeletal muscle cells (Figure 5G-H). Using ImageJ ROI tools, we were able to isolate single muscle cells and record calcium transients (Figure 5I). These data combined provide robust support for the use of MCA in a wide range of functional imaging applications across model organisms and tissue types.

## Discussion

Fluorescent imaging strategies have emerged as a fundamental experimental technique for understanding functional dynamics by capturing the activity of cells.^64^ The utility of calcium imaging has been particularly impactful for studying the brain and neural circuit function. In the brain, many functional imaging strategies often record from hundreds if not thousands of neurons, which results in technically complex datasets requiring sophisticated computation. Moreover, analyzing calcium imaging data is further complicated due to the diversity of file formats, proprietary software limitations, and the need for programming knowledge.^24,65^ Considering these hurdles, many research groups develop custom software solutions.^16,66^ However, limitations of such custom approaches are that these boutique packages may be challenging to broadly apply by other groups as they require advanced coding or proprietary software.^12,13^ To address these challenges, we developed the MCA (MultiCellular Analysis) toolkit for ImageJ, a widely used open-source image processing software. Our goal is to provide a streamlined, flexible, and user-friendly tool that enables researchers to efficiently process and analyze functional imaging data.

Our MCA plugin has addressed many challenges from other functional imaging analysis solutions and offers a user friendly, widely applicable, and easily accessible alternative. A major contributor to the increased accessibility of the MCA toolkit is that it is built within ImageJ, which has been an industry standard image analysis platform for over 25 years^17,18,67^. We demonstrate that our plugin has a variety of functionalities applicable to calcium imaging analyses such as motion correction, cell detection, data conversion, annotation, and activity prediction. Many of these methods have been developed in ImageJ previously yet not organized to work together in one application intended for functional imaging. Moreover, MCA has the flexibility to leverage all other ImageJ functions and vast repertoire of plugins for image analysis.

While there are a variety of established tools for functional imaging analysis such as Caiman or Suite2p, these tools require programming knowledge (Caiman) or limit extended functionality from the program (Suite2p)^12,13^. These tools alone for functional imaging analysis have demonstrated themselves to be powerful tools designed for very specific purposes^2,68–71^. We wanted to allow functional imaging analysis to be completed alongside other analytical techniques to reduce the necessity of employing multiple software required for each unique experiment. Therefore, completing functional imaging analysis in ImageJ while having access to other processes in ImageJ streamlines analysis of scientific imaging data.

## Limitations of the study

A primary goal of MCA was to provide unified software for completing routine analyses. As such, the choice was made to design this program within ImageJ compared to developing it within other languages such as Python or MATLAB. This poses a future challenge as many analytical techniques are not natively developed using Java, although Java remains one of the most popular programming languages today. Many newer image analysis-based applications are developed with Python which sometimes aren’t readily portable to an ImageJ plugin. However, ImageJ/Fiji have been addressing this as they continue to expand resources for different development suites within ImageJ providing methods for designing macros in a variety of languages which can be incorporated with plugins. The ability to easily add or extend functionality into MCA through Java based methods or ImageJ macro-based methods highlights the ability for MCA to remain a modular and user-friendly application for functional imaging analyses. MCA provides a powerful, user-friendly option for analyzing functional imaging data in ImageJ. However, MCA is designed to primarily focus on single plane recordings and currently is not readily adaptable to volumetric imaging. In addition, MCA is oriented at providing visual output for nearly all functions available and generating images to be displayed is inherently computationally expensive. Other software that provides visual output optionally, such as Caiman or Suite2p, benefit from increased processing speed through analyses.

## Methods

### Animal Husbandry

All usage of animal and experiments were approved by the West Virginia University Institutional Animal Case and Use Committee. <u>Zebrafish</u>: All experiments utilized the Tupfel long-fin wildtype genetic background. All functional imaging experiments took place at or prior to 7 days post fertilization. Larvae were raised at 28.5°C in a 14/10 hour light/dark cycle prior to use. Larvae were raised in E3h embryo media. Transgenic lines used were *Tg(y279:Gal4)*, *Tg(elavl3:H2B-GCaMP6f), Tg(UAS:nsfb-mCherry), and Tg(y264:Gal4)*. *<u>Drosophila</u>*: All fly stocks were raised on a standard cornmeal/agar/yeast medium at 24°C on a 12:12 light/dark cycle at ∼60% humidity. Male and female, 3-5 day old flies were used for recordings. Transgenic lines used were w1118; ;UAS-GCaMP7f/+;R32F10-Gal4/UAS-GCaMP7f. <u>Mice and mouse welfare</u>: *Cntnap2*^−/−^ mice (https://www.jax.org/strain/017482, stock number 017482, The Jackson Laboratory) were bred as heterozygous (HET) pairs, resulting in *Cntnap2*^+/+^ (wild type, WT), *Cntnap2*^−/−^ (KO), and *Cntnap2*^+/−^ (HET) offspring. *Cntnap2* KO mice and WT littermate controls of either sex were used beginning at eight weeks of age. Data presented is from a confirmed wildtype animal. Animals were housed in a vivarium with *ad libitum* access to food and water, on a 12/12 hr light/dark cycle. All animal procedures were performed in accordance with the Hussman Institute for Autism animal care committee’s regulations and those of the University of Maryland Baltimore IACUC where recordings were performed.

### MCA development

The MCA plugin was developed to utilize the IJ 1.54f library. MCA was designed in IntelliJ and is built with Maven3 and compiled for Java 8. All UI components were developed using Javax or the abstract widget toolkit (AWT). Motion correction algorithm was based on the “Template_matching” plugin developed for ImageJ and used the OpenCV library for the template matching function^23,72^. Annotation tools such as the “Point grouping” and “Polygon grouping” functions utilized tools available through the toolbar and ROI manager in ImageJ. The Cell Manager UI framework was based on the native ROI Manager in ImageJ. ROI signal was extracted from each ROI using the average pixel value within each ROI for every frame and exported using the “ResultsTable” class available in the IJ 1.54f library. Single peaks were determined using a sliding window method described in ^73–76^. This method takes the standard deviation of the window size for every subsequent frame beyond the size of the window and adjusts the window based on an influence factor of the signal set by the user. Then with a set threshold of standard deviation, points falling outside the threshold are considered “1” for a significant increase in signal, or “-1” for a significant decrease in signal, or “0” for no change in signal. Signal data was filtered using a gaussian smoothing filter described in ^77^. The model for Cellpose was trained using 117 sum projection images of the zebrafish thalamic region expressing *Tg(elavl3:H2B-GCaMP6f)* using Cellpose’s built in training method.

#### Visual calcium imaging

Visual stimulus calcium imaging data was used from Hageter et. al 2023^31^. All individuals from the prior publication were reanalyzed. Cellular maps containing *Tg(elavl3:H2b-GCaMP6f)* and *Tg(y279:Gal4)* were used for cellular annotation. Cell ROIs were generated using a custom trained Cellpose model (H2BGCaMP), a binary threshold was applied, and segmented using a watershed segmentation algorithm^78^. These were then grouped together based on anatomical landmarks into 3 regions in the thalamus. The greater thalamus, the lateral thalamus, or the posterior tuberculum. Cells expressing *Tg(y279:Gal4; UAS:nsfb-mCherry)* were point labeled. All images had a rolling ball radius of 20 pixels of background subtraction prior to conversion to 32-bit ΔF/F format. Each fish had two acquisitions which were averaged together prior to exporting. All data was smoothed with a gaussian filter. Peaks were detected using a lagging window of 30 frames, and a threshold of 3 times the standard deviation of the window for each subsequent timepoint following the width of the window.

#### Acoustic calcium imaging

Acoustic stimulus calcium imaging data was collected for this manuscript. Larvae were housed under normal conditions and raised in 200 µM PTU until 5-6 dpf where they were anesthetized using tricane and embedded in 2% LMP agarose. Larvae were acclimated to the imaging environment for 5 minutes prior to recording. Larvae were imaged on a Scientifica Vivoscope two-photon with a 16x water immersion objective, and Spectra-Physics MaiTai laser tuned to 940nm. Acoustic stimulation was provided using a skinny mini excited 9mm, 1W, 4Ohm (DaytonAudio).^79^ The speaker was delivered by a 2×15W class D audio amplifier board (DaytonAudio). Stimulus was controlled through a custom script written in IDL eventtimer^80^. Generated stimulus consisted of incrementally increasing voltage sent through the amplifier to deliver a 300 msec acoustic stimulus with 10 msec of ramp. Stimulus were generated with 1000 samples and consisted of 1000 Hz. Larvae were imaged for two repeats of 5 auditory stimulus separated by 30 seconds.

#### Fly calcium imaging

Drosophila calcium imaging was completed on a Scientifica Vivoscope two-photon microscope with a Spectra-Physics MaiTai sapphire laser at 940 nm wavelength. Imaging was conducted on flies expressing GCaMP7f under the control of the R32-GAL4 driver line. Images were acquired using ScanImage acquisition software for MATLAB. Images were acquired at 3.4 Hz. Flies were prepared by anaesthetizing on ice and fixed with LED-UV plastic welder (BONDIC, SK8042, NY). ACV was perfused onto the antennae from a 1:100 dilution in distilled water and odor stimuli were delivered as a 1-second puff directed at the antennae using a custom built odor deliver system.^81^ ROIs were manually drawn with native ImageJ ROI tools to manually define unique glomeruli. These ROIs were synced to MCA and imaging series were converted to ΔF/F format using the “Convert to DF/F” in MCA with a set baseline of 10 frames. Peaking activity was determined using a sliding window of 5 frames and golemuli with >3 standard deviations of change in signal for at least 3 frames following ACV perfusion were considered significant responses.

#### Mouse calcium imaging

Mouse calcium imaging was completed on a Zeiss LSM-780 microscope tuned to 940 nm with 20x immersion objective using a head fixed wildtype mouse expressing GCaMP6s and 3 different fields of view of the superficial primary visual cortex (V1). Images were acquired using ZEN Black software at a frame rate of 4.2 Hz. Stimulus presented was a randomly displayed set of full-field drifting sinusoidal gratings at 100% contrast, with temporal frequency of 1 Hz, and with a spatial frequency of 0.02 series cycle per degree. The stimulus presented cycled pseudo-randomly through 8 different angular orientations each 45 degrees apart, and the sequence was repeated ten times. Visual stimuli were generated in MATLAB using PsychoPhysicsToolbox, and presented on a gamma-corrected computer monitor (32” display) placed 20 cm distant from the animal. Visual stimulation was presented for 4 seconds at a time separated by 4 seconds of a blank grey screen. Imaging files were imported into ImageJ and cells were manually labeled from a sum projection of the recording series and imported into the ImageJ ROI manager. From the ROI manager, ROIs were synced to the Cell Manager and image stacks were converted to ΔF/F format using a set baseline of 20 frames. Peaking activity was determined from a sliding window of 10 frames and cells with ±3 standard deviations at the timepoint following the sliding window were considered significant increases or decreases in cell activity, respectively.

#### Muscle calcium imaging

Calcium imaging of the zebrafish skeletal muscle was completed using wildtype (TL) zebrafish injected with *Tg(actc1b:GCaMP5)* plasmid acquired from Addgene. Zebrafish were injected at the single cell stage with 4.6 nL of plasmid at a concentration of 25 ng/L diluted in 1x Evans solution. Zebrafish were raised in normal housing conditions with media replaced daily and sorted on an epifluorescent dissecting microscope at 5 dpf. Zebrafish were anaesthetized with 0.1% Tricaine MS-222 for two minutes prior to being mounted laterally in 1% low melting point agarose (LMP). Mounted zebrafish were stored at 28°C for 10 minutes prior to imaging. Imaging was completed on the same multiphoton microscope described in other zebrafish imaging experiments. Zebrafish were recorded for two 3 minutes at 1 Hz under 690 nm LED illumination with no further stimulation provided. Recordings were loaded into ImageJ and concatenated in sequence. ROIs were manually drawn around unique muscle cells. We used a baseline period of 60 frames to calculate ΔF/F using MCA’s “Convert stack to DF/F” function.

#### Data analysis

All statistical analysis was completed in R using package rstatix^82,83^. All t-tests are two tailed. Significance was determined based on an alpha value of 0.05. Data presented represents the mean ± standard error of the mean.

## Data and code availability

All original data was uploaded to Mendeley and can be found at <Mendeley link posted with peer reviewed publication>. Previously published, publicly available data used in this manuscript can be found using accession numbers listed in the key resources table. All original code used in the development of MCA is available at https://github.com/JohnHageter/Multi-Cell-Analysis.

## Acknowledgements

We want to thank Kevin Daly, Jeff Mumm, and Sadie Bergeron for helpful insights through the development of this work. We would also like to thank Curtis Reuden and colleagues for the maintenance and future development of Fiji and ImageJ. This work was supported by National Science Foundation cooperative agreement OIA2242771, National Eye Institute R15EY036226, and National Institute of General Medicine P20GM144230 awarded to Eric Horstick; NIH DC-016293, NSF IOS 2114775, and AFOSR DURIP awards FA9550-19-1-0179 and FA9550-20-1-0098 awarded to Andrew M. Dacks; and Hussman Foundation grant HIAS18001 and National Science Foundation (NSF) Established Program to Stimulate Competitive Research (EPSCoR) Track-1, Award OIA-2242771 awarded to M. Bridi.

## Author contributions

E.J.H and J. Hageter conceived the project and wrote the manuscript. J. Hageter developed software, analyzed data, performed experiments. J.J., M.B., performed experiments. A. DelGaudio, M.L., B.J., T.R., J. Holcomb, A.F.T, and J.J. analyzed data. M.B., A.M. Dacks, and E.J.H acquired funding.

---

**Supplemental figure 1:**
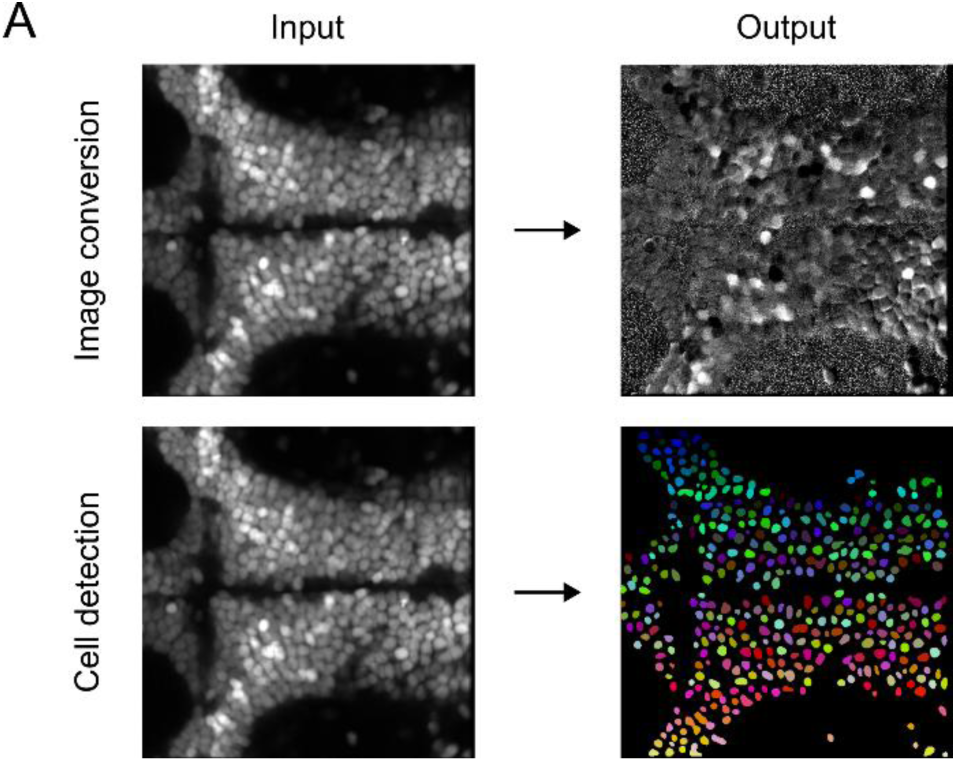
MCA functions have visual output. **A.** Representative input image and output image for MCA’s image conversion function (top) or Cellpose mediated cell detection (bottom)

**Supplemental Figure 2:**
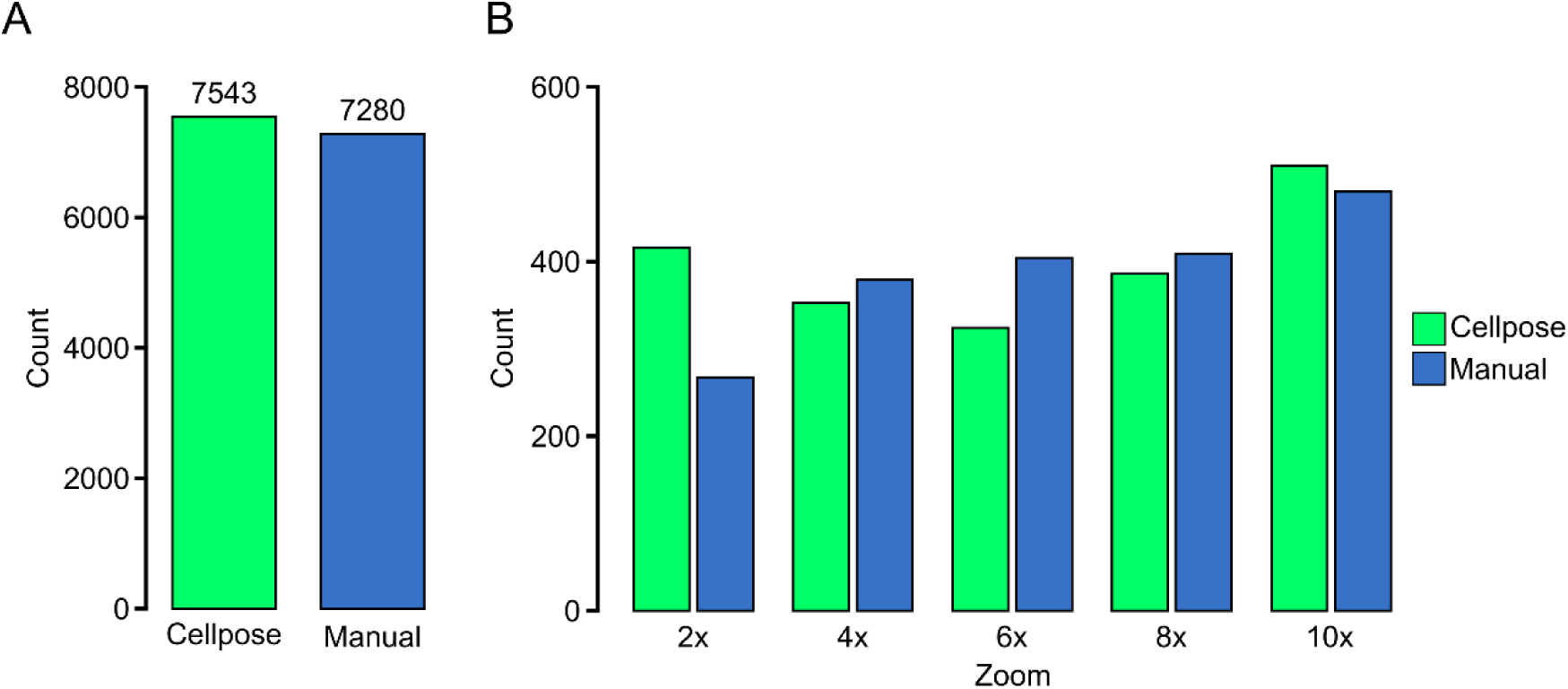
Cellpose model verification. **A.** Counts of cells in the thalamus extracted from the trained H2B-GCaMP Cellpose model (green, N=19), or manually counted cells (blue, N=19). The number above bar represents the total number of cells counted among all larvae. **B.** Comparison of cell counts from Cellpose output, or manual counting based on digital zoom from a 20x water immersion objective.

**Supplemental Figure 3:**
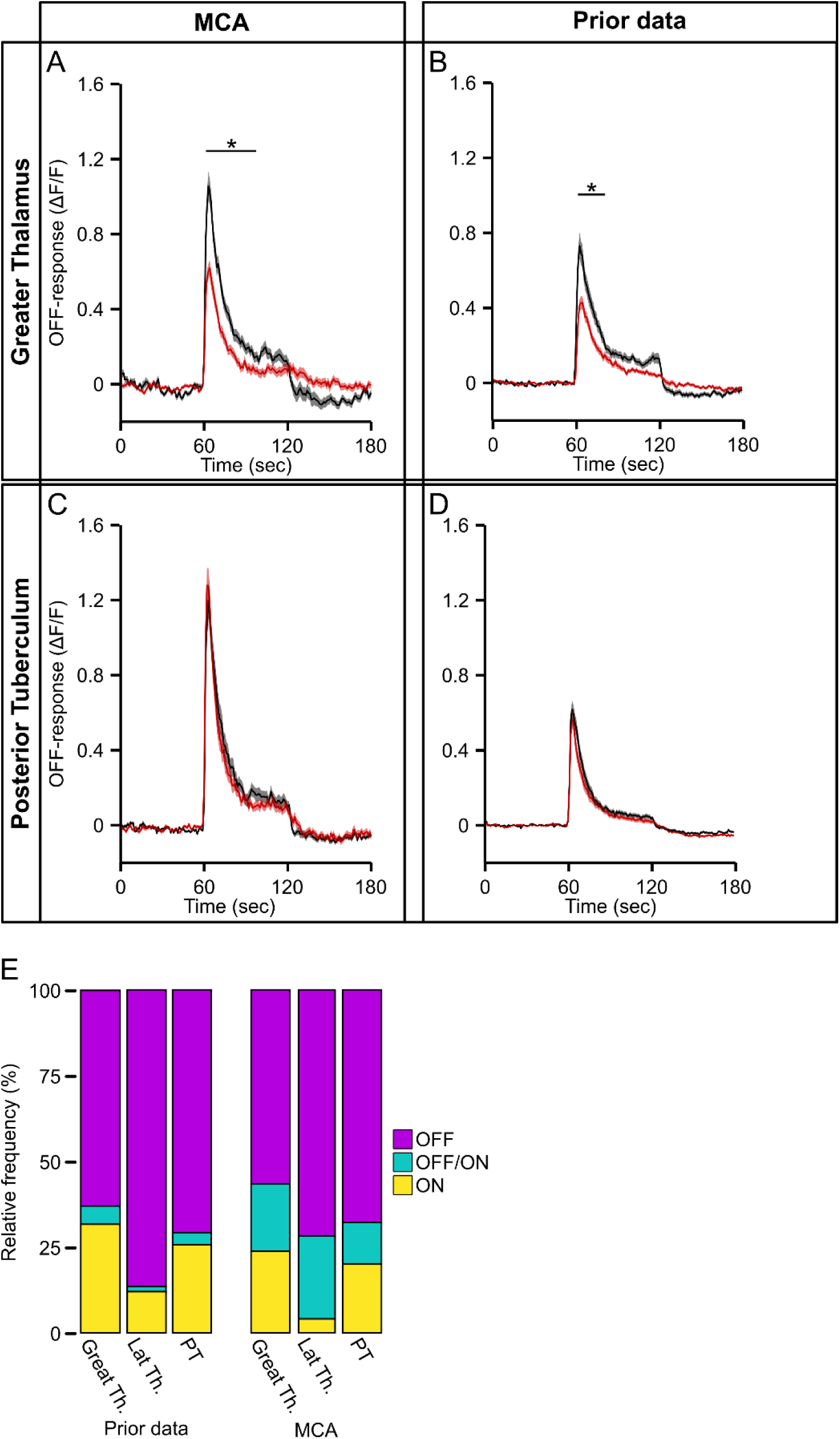
MCA data export validation. A-B. Average response for OFF cells in the Greater Thalamus analyzed with **A.** MCA or **B.** previously published for matched (black) and opposed (red) hemispheres. (MCA: Matched n=84, opposed n=80; Prior data: Matched n=63, opposed n=80). **C-D.** Same as in **A-B.** for the Posterior Tuberculum analyzed with **C.** MCA or **D.** previously published (MCA: Matched n=123, opposed n=127; Prior data: Matched n=170, opposed n=148). **E.** Relative frequency of cellular response types from each region excluding non-responding cells for OFF (magenta), ON (yellow), or OFF/ON (cyan) responsive cells. * indicates p<0.05 two tailed t-test between hemispheres for at least 10 consecutive timepoints.

